# Role of lipid packing defects in determining membrane interactions of antimicrobial polymers

**DOI:** 10.1101/2022.06.08.495334

**Authors:** Samapan Sikdar, Garima Rani, Satyavani Vemparala

**Author notes:** Equal Contribution (Samapan Sikdar, Garima Rani).

## Abstract

Understanding the emergence and role of lipid packing defects in the detection and subsequent partitioning of antimicrobial agents into bacterial membranes is essential for gaining insights into general antimicrobial mechanisms. Herein, using methacrylate polymers as a model platform, we investigate the effects of inclusion of various functional groups in the biomimetic antimicrobial polymer design on the aspects of lipid packing defects in model bacterial membranes. Two antimicrobial polymers are considered: ternary polymers composed of cationic, hydrophobic and polar moieties and binary polymers with only cationic and hydrophobic moieties. We find that differing modes of insertion of these two polymers lead to different packing defects in the bacterial membrane. While insertion of both binary and ternary polymers leads to an enhanced number of deep defects in the upper leaflet, shallow defects are moderately enhanced upon interaction with ternary polymers only. We provide conclusive evidence that insertion of antimicrobial polymers in bacterial membrane is preceded by sensing of interfacial lipid packing defects. Our simulation results show that the hydrophobic groups are inserted at a single co-localized deep defect site for both binary and ternary polymers. However, the presence of polar groups in the ternary polymers use the shallow defects close to the lipid-water interface, in addition, to insert into the membrane, which leads to a more folded conformation of the ternary polymer in the membrane environment, and hence a different membrane partitioning mechanism compared to the binary polymer, which acquires an amphiphilic conformation.

## 1. Introduction

Exploration of clinical usage of antimicrobial peptides and their synthetic mimics has been at the forefront in last few years in the wake of emergence of antibiotic resistance among pathogens [1, 2, 3, 4, 5, 6]. In particular, considerable research efforts have been put in to developing biomimetic polymers as a viable alternative to naturally occurring antimicrobial peptides which suffer from issues including proteolysis, bioavailability and poor *in-vivo* activity [7, 8, 9, 10, 11]. The primary goal in design of these antimicrobial polymers is to capture the essential antimicrobial properties and not necessarily replicate the exact structural details. Towards this end, many synthetic polymers comprising of the critical functional groups, hydrophobic and cationic, have been studied[12, 13, 14, 15]. The primary role of the cationic groups is to differentiate between largely zwitterionic host cell membranes and anionic bacterial membranes, while the hydrophobic groups play crucial role in membrane disruption. The presence of these two functional groups has also been implicated in facially amphiphilic structures of linear antimicrobial agents. However, an inspection of the available databases on antimicrobial peptides strongly suggests the presence of other functional groups like polar and negatively charged residues in significant number of antimicrobial peptides [16]. It is important to understand how differently they interact with bacterial membranes when compared to the previously described binary composition in designing better antimicrobial mimics. Recently we showed that replacing some of the hydrophobic groups in a binary biomimetic polymer with polar groups significantly alters how the resultant ternary polymers interact in solution phase and with bacterial membranes[17, 18]. The studies also strongly suggest that the inclusion of polar groups smears the overall hydrophobicity of such polymers and result in a globular conformations when partitioned into the bacterial membrane, likely mimic of defensin-like antimicrobial peptides [19, 20].

A crucial factor having direct influence on AM agent - membrane interactions is the lipid composition of the membranes. Changing the lipid composition of membranes can be broadly classified into changes in (a) lipid head groups, and (b) lipid tails. In the context of AM agent - membrane interactions, the two types of changes can be envisaged to predominantly influence different stages in the antimicrobial action. The recruitment of AM agent molecules involves the recognition through long - range electrostatic interactions, followed by adsorption and partitioning of the agent molecule. The latter involves specific interactions with lipid head group atoms such as hydrogen bonding, in addition to electrostatic and vdW interactions.The interfacial region of a lipid bilayer is highly dynamic, and the sub-nanometer structure of the lipid bilayer-water interface is subject to constant dynamical fluctuations, often resulting in transient unfavorable exposure of hydrophobic lipid tail group atoms to interfacial water. Regions of hydrophobic exposure are defined as interfacial lipid packing defects, and they provide favorable binding sites to the hydrophobic groups of molecules adsorbed at the lipid bilayer interface. The interactions of unpartitioned agent molecules with the membranes is thus essentially an interfacial phenomenon having greater dependence on the head group composition, compared to the composition of lipid tails. Partitioned agent molecules, in contrast, interact extensively with the lipid hydrophobic tails. Changes in lipid tail composition have a strong influence on properties such as bilayer thickness and order in lipid tail [21]. Alteration of such properties can critically alter the conformations of partitioned agent molecules, as well as their influence on the bilayer properties.

Considerable recent research efforts have been focused on the specific role of membrane structure in aiding the recognition and subsequent partitioning of the membrane active agents into the membranes. The large heterogeniety that exists in the different membranes in cells or organelles can be directly related to the composition of the lipid molecules in such membranes. The main differences among the lipid molecules include overall charge on the head group, size of the head group, saturation along the lipid tails. These determine both how a membrane active agent may perceive a membrane patch and also its interaction and possible partitioning into such membranes. Recently there has been much focus on the non-ideal packing of lipid head groups leading to interfacial packing defects, which has also been suggested to play an important role in the partitioning of amphiphilic agent molecules to the membrane interior [22, 23]. There are several examples of proteins and peptides which can sense these interfacial packing defects, insert and subsequently partition into them [24, 25, 26, 27, 28]. Recently, it has been reported that the recruitment of amphiphilic molecules by lipid bilayers is modulated by the topography of membrane interfacial region [22]. At the core of the notion lies the concept of interfacial lipid packing defects, which results in the transient exposure of hydrophobic lipid tail atoms to hydrating water outside the membrane. Interfacial lipid packing defects can be categorized into chemical and geometrical defects, based on the relative depth of the defect sites with respect to the nearest glycerol backbone. By definition, all lipid packing defects resulting in hydrophobic exposure are chemical defects. If the defects involve exposure of hydrophobic tails located deeper than the position of nearest glycerol atoms, they are further classified as geometrical defects [23]. It has been suggested that membrane active molecules are intrinsically capable of detecting such defects, and favorably bind to them [22]. The presence of conical lipids with relatively small head groups compared to lipid tail cross section, such as PE groups with intrinsic negative curvature, have been known to result in enhancement of such interfacial lipid packing defects [23]. In comparison, the effective shapes of lipid molecules with PC head groups are cylindrical, allowing them to pack more effectively in the planar bilayer phase.

In this paper, we investigate how the presence of different functional groups in the antimicrobial polymers sense, affects, exploits the lipid packing defects present in the model bacterial membranes for partitioning into the same. To this end, we compare the partitioning dynamics of ternary polymer composed of charged cationic, hydrophobic and polar groups with that of binary polymer, lacking the polar residues. The action of antimicrobial polymers enhances both the number and the size of lipid packing defect sites. The hydrophobic residues of both polymers sense the interfacial lipid packing defects and occupy co-localized deep defect sites, thereby resulting in partitioning of these polymers into membrane interior. On other hand, the polar residues of ternary polymer participating in extensive hydrogen bonding network with lipid headgroups, tend to reside in shallow defect sites close to the bilayer water interface. This tendency of hydrophobic and polar residues to occupy defect sites of varying depths leads to a more folded conformation of ternary polymer in contrast to the facially amphiphilic linear conformation of binary polymer, resulting in different partitioning mechanisms. That these antimicrobial polymers can also sense membrane topography is indicative of a generalized role of interfacial lipid packing defects in governing partitioning of different membrane active agents.

**Table 1:** Proportion and sequence of AEMA (A), HEMA (H) and EMA (E) monomers in the random polymer models (T and B). In all the model polymers, degree of polymerization (DP) = 19 and the number of cationic side chain groups are fixed to be 6 per polymer.

| Model copolymers | DP | Group proportion (AEMA, HEMA, EMA) | Group Sequence |
| --- | --- | --- | --- |
| Ternary (T) | 19 | +6, 5, 8 | E-H-A-E-H-A-H-E-A-E-E-A-E-H-H-A-E-E-A |
| Binary (B) | 19 | +6, 0, 13 | E-E-A-E-E-A-E-E-A-E-E-A-E-E-E-A-E-E-A |

## 2. Models and methods

Atomistic MD simulations were performed on systems containing binary (referred to as “model B”) and ternary methacrylate (referred to as “model T”) polymers with model membranes in the presence of water and salt ions. The binary polymer consists of two functional groups, cationic ammonium (AEMA), hydrophobic alkyl (EMA) and the ternary polymer has an additional neutral hydroxyl (HEMA) group. Chemical structures of EMA, AEMA and HEMA are given in Fig.1A. Both ternary and binary polymers have a degree of polymerisation (DP) = 19 with the groups in a random sequence with a composition of 6 AEMA units, 8 EMA units, 5 HEMA units for model T polymers and 6 AEMA units, 13 EMA units for model B polymers. For further details and an in depth study of such polymers and their conformations in solution and membrane phase we refer to our previous works [17, 18]. The bacterial membrane consists of 38 POPG and 90 POPE lipid molecules per leaflet to mimic the inner membrane of the Gram negative bacteria [29]. CHARMM-GUI’s Membrane Builder module [30] was used to construct the the membrane system, similar to our previous works [31, 32, 18]. All simulations were performed using the NAMD simulation package [33]. The polymer-membrane systems were simulated in the isothermal-isobaric ensemble (P = 1 atm and T = 310 K) with periodic boundary conditions and a time step of 2 fs. CHARMM 36 forcefield was used for the membrane systems [34], and the force field for polymers was from our previous published simulation studies [35, 36, 31]. The model T polymer-membrane system was simulated for 900 ns and the model B polymer-membrane system was simulated for 700 ns (for more details of the simulation study of such systems, we refer to our previous work [18]).

**Figure 1:**
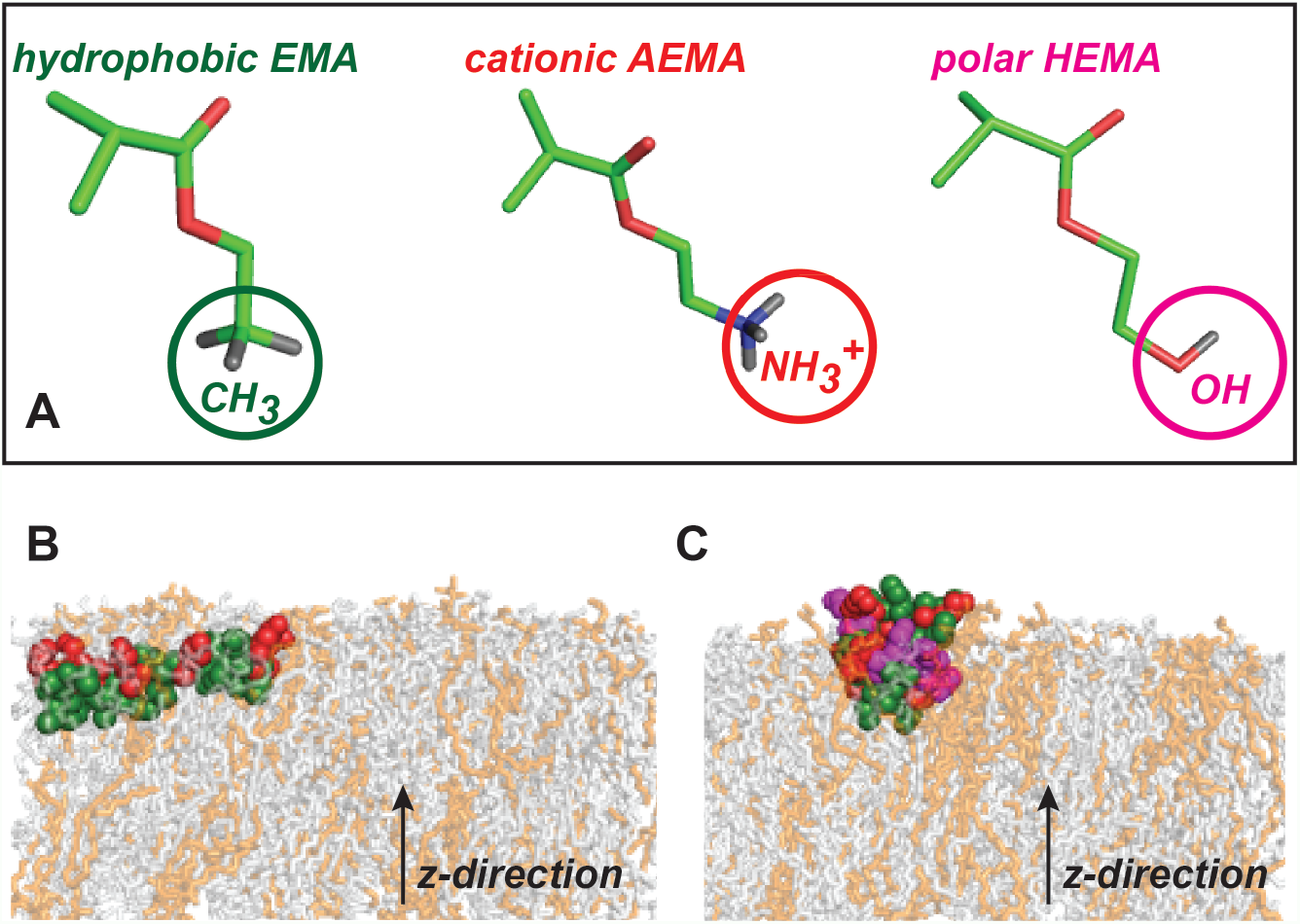
(A) Chemical structures of EMA, AEMA and HEMA groups considered in the model polymers. The final snapshots of Binary (B) and Ternary (C) polymers interacting with model bacterial membrane, composed of POPE (gray) and POPG (orange) lipids. The monomeric units: cationic AEMA, hydrophobic EMA and polar HEMA of the two polymers are shown in red, green and magenta, respectively. For more details of polymer-membrane interactions, please see Ref.[18].

The analysis of the membrane-polymer systems is performed using Visual Molecular Dynamics (VMD) [37]. Further analysis pertaining to detection of lipid packing defects is performed using Packmem[38] software and in-house Fortran codes. The Packmem algorithm divides the bilayer x-y plane into 1 Å *X* 1 Å grids. It then scans along the membrane normal (z-direction) from bilayer-solvent interface down to 1 Å below the average level of C2 atoms of lipid glycerol moieties to identify voids resembling packing defect sites. These defect sites are qualitatively characterized into deep or shallow depending on their relative depth with respect to the C2 atom. Further quantification is done based on the area (*A*) of defect site. The calculation is repeated for each frame to compute the number, area and location of defect sites in a given frame.

## 3. Results

The earlier study demonstrated significant differences in both conformation and binding modes of ternary (model T) and binary (model B) polymers in their interactions with model bacterial membrane [18] (Figures 1B and 1C). While the binary polymer acquired a linear facially amphiphilic conformation upon insertion, the ternary polymer adopted a more folded conformation, aligned in the direction of membrane normal. The inclusion of polar HEMA units in addition to cationic AEMA and hydrophobic EMA monomeric units resulted in deeper partitioning of ternary polymer compared to binary case, the latter comprising of only cationic and hydrophobic moieties [18]. In the present work, we elucidate how the interfacial topography of bacterial membrane contributes to the observed differences in binding modes of these polymers. The interfacial packing defects are computed over each frame of the MD trajectories using Packmem[38]. We first estimate the abundance of defect sites of varying sizes by drawing a distribution of the same over the last 500 ns of equilibrated trajectories corresponding to each polymer-membrane system.

### 3.1 Distribution of defect size

The distribution of defect size, *P* (*A*) (shown in semi-log scale, Fig. 2), where *A* represents the area of individual defect sites, is computed for the two leaflets separately. Both model B and model T polymers reside near the upper leaflet, denoted as *L*0, while *L*1 represents the lower leaflet. The *P* (*A*) of deep defects in presence of binary (Fig. 2A) and ternary (Fig. 2B) polymers reflect relative population of different defect sizes in the two leaflets. We observe distinctly different deep defect size distributions in these leaflets. The distal leaflet, *L*1, is characterized by presence of small defect sites, within 50 Å^2^. However, in presence of either polymers, the proximal, *L*0 leaflet not only exhibits enhanced probability of similar small sized defects but also possesses significantly large defect sites 100 - 150 Å^2^. Thus the distal leaflet seemingly remains unperturbed by the action of polymers, while both number and size of deep defects increase in the proximal leaflet. Owing to the presence of these large defects in *L*0, the corresponding deep defect *P* (*A*) exhibits multi-exponential behaviour in contrast to the single exponential decay profile observed in *L*1. This deviation from single exponential behaviour in the limit of large defects is in agreement to earlier studies [26, 27].

**Figure 2:**
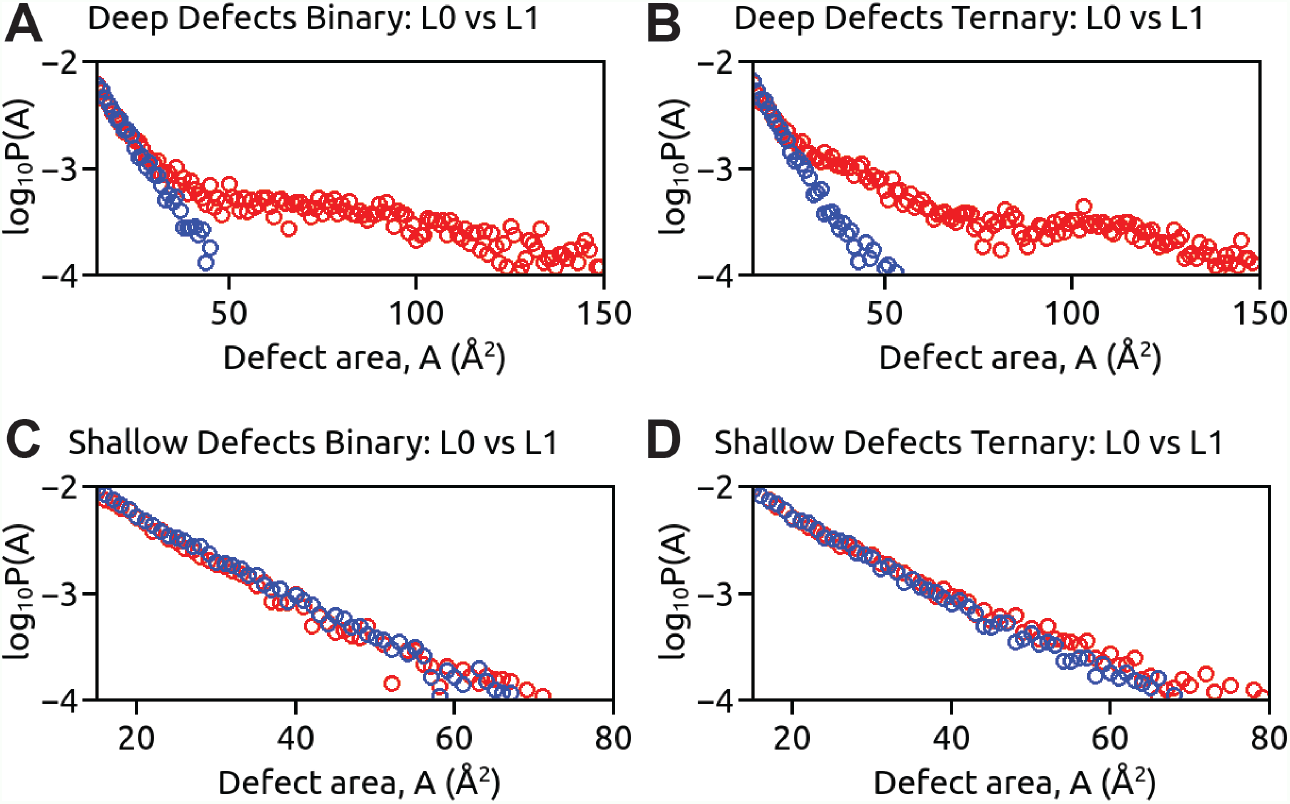
The size distributions, *log*_10_*P* (*A*), where *A* represents the area of individual defect sites are illustrated for deep lipid packing defects populating the proximal, *L*0 (red) and distal *L*1 (blue) leaflets upon interaction with (A) Binary and (B) Ternary polymers. The size distributions of shallow lipid packing defects in the two leaflets are shown for (C) Binary and (D) Ternary polymers.

On the other hand, the size distribution, *P* (*A*) of shallow lipid packing defects remains almost similar in both leaflets upon insertion of model B polymer (Fig. 2C). However, a modest increase in shallow defects is observed in the proximal *L*0 leaflet compared to *L*1 due to interaction with the ternary polymer (Fig. 2D), as discussed in the following section. A comparison between the two defect types demonstrate that deep defects are more prevalent than shallow defects in the proximal leaflet. The abundance of large deep defect sites induced by action of these polymers may facilitate their partitioning into the bacterial membranes. As such, several studies[24, 25, 26, 27, 28] have confirmed the role of these lipid packing defects as transient binding “hotspots” for peptide insertion and subsequent partitioning. In the following section we show how these polymers can also sense lipid packing defects similar to proteins and peptides and consequently describe their mechanism of partitioning into the bacterial membrane.

### 3.2 Sensing Lipid Packing Defects

In order to elucidate the role of deep lipid packing defects in driving partitioning of the polymers into bacterial membrane, we study the insertion dynamics of a representative set of hydrophobic EMA monomer units (residues) within the model B polymer. We follow the insertion dynamics of these residues by calculating the individual distance (z-distance) of residue centre of mass from the average level of C2 atoms of lipid molecules. A negative value of z-distance corresponds to residue insertion below the average C2 level. Simultaneously, we scan the interface region to track appearance of any underlying deep lipid packing defect co-localized with these residues. We identify a single large colocalized deep defect beneath these residues. Fig. 3A illustrates the insertion dynamics of the EMA residues and the appearance of a single large co-localized defect. Within 50 ns of simulation run, a co-localized deep defect of area around 50 Å^2^ appears, facilitating insertion of one of the EMA residues, following which the area of defect site increases. Then close to 170 ns, with the co-localized deep defect area fluctuating around 100 Å^2^, the other representative EMA residue gains entry into the membrane milieu. Owing to these discrete events of hydrophobic insertions, the co-localized deep defect first increases in size and then gets stabilized, driving entry and subsequent partitioning of model B polymer into the bacterial membrane.

**Figure 3:**
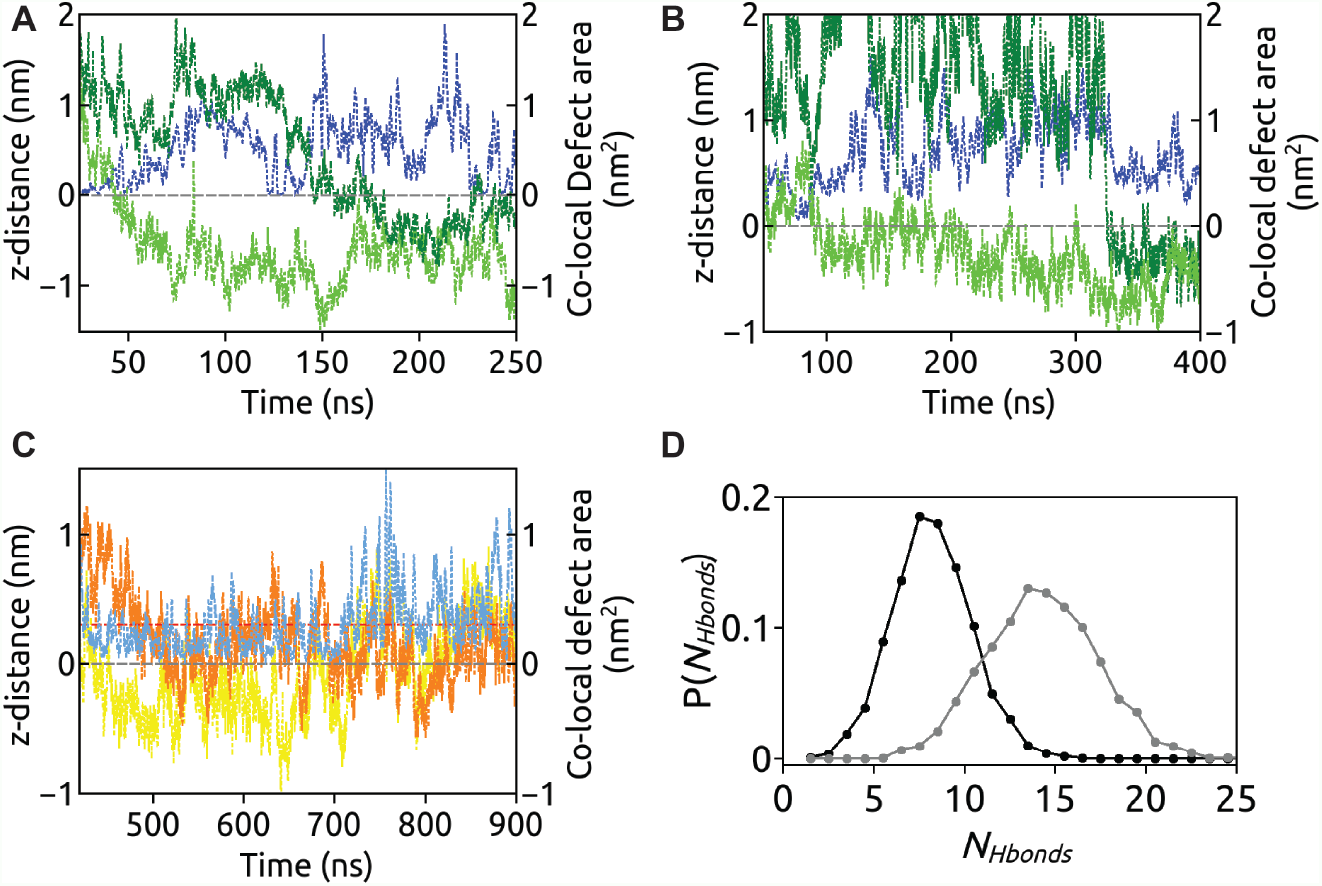
The insertion dynamics of two representative hydrophobic EMA residues (shown in different shades of green) belonging to Binary (A) and Ternary (B) polymers into a colocalized deep defect site (blue). The insertion dynamics of two representative polar hydroxyl HEMA residues (in yellow and orange) of Ternary (C) polymer exhibiting intermittent residence in a co-localized shallow defect (light blue) between the average levels of C2 (grey) and phosphate (red) atoms of lipid molecules. (D) The distributions of number of hydrogen bonds, *P* (*N*_*Hbonds*_) indicate that the Ternary polymer (grey) exhibits strong hydrogen bonding network with surrounding lipid and solvent molecules compared to the Binary (black) polymer.

On a similar note, we also probe the partitioning dynamics of model T polymer considering two representative hydrophobic EMA residues (Fig. 3B). The appearance of a co-localized deep defect drives hydrophobic insertion near 100 ns. Following this insertion event, the deep defect steadily increases from 50 Å^2^ to nearly 100 Å^2^, promoting another hydrophobic entry at 350 ns. It is worthwhile to note that a shallow defect site also appears in close proximity to the deep defect, albeit smaller in size 30 Å^2^. The shallow defects are characterized by depths above the level of C2 atoms of glycerol moieties of lipid molecules. We show the existence of this shallow defect intermittently occupied by some representative polar residues in Fig. 3C. Being polar, these residues find favourable to reside close to the lipid-water interface, between the levels of C2 atoms and phosphate (P) atoms of lipid molecules, representing shallow insertion.

These polar hydroxyl groups in ternary polymer can act both as donor and acceptor of hydrogen bonds, resulting in extensive hydrogen bonding network with lipid headgroups and interfacial water molecules. We observe significant differences in the hydrogen bonding pattern of the two polymers with surrounding solvent and lipid molecules. The distribution of number of hydrogen bonds, *P* (*N*_*Hbonds*_), shown in Fig. 3D indicates strong tendency of ternary polymer to form hydrogen bonds, *< N*_*Hbonds*_ *>*∼ 15. However, the absence of polar groups in binary polymer leads to reduced occurrence of hydrogen bonds with the lipidwater interface (*< N*_*Hbonds*_ *>*∼ 7). This in particular explains the intermittent co-localization of polar residues leading to the modest enhancement of shallow defects in *L*0 (Fig. 2D) upon interaction with ternary polymer but not with binary polymer.

The final MD snapshots of membrane embedded binary and ternary polymers superposed with the co-localized deep and shallow defect sites are illustrated in Fig. 4A and 4B, respectively. Having studied the co-localization of defect sites with polymers, it is worthwhile to investigate if these defect sites also co-localize with a particular lipid type. The model bacterial membrane considered in the current study being composed of POPE and POPG lipid molecules, we determine if there exists a preferential sorting of lipids towards defect sites in the following section.

**Figure 4:**
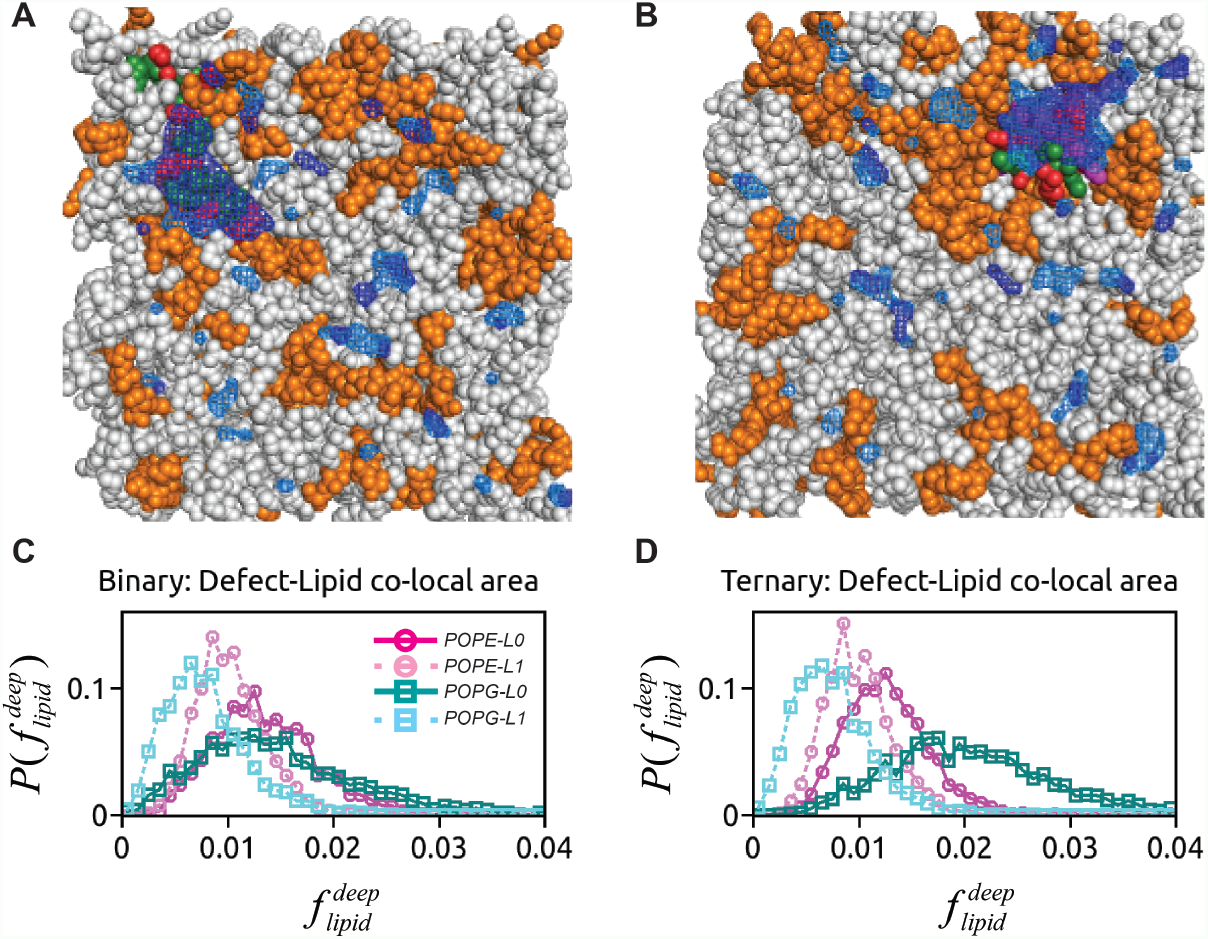
Illustrations of the final snapshots of Binary (A) and Ternary (B) polymers completely embedded into the large co-localized deep defect (dark blue) surrounded by shallow defects (light blue). The POPE and POPG molecules are shown in gray and orange, respectively. The clustering of POPG molecules is observed in vicinity of the Ternary polymer (B) insertion site. The monomeric units: cationic AEMA, hydrophobic EMA and polar HEMA of the two polymers are shown in red, green and magenta, respectively. The distributions of fraction of lipid surface areas co-localized with deep defects in each leaflet, 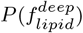 corresponding to two lipid types: POPE and POPG are illustrated for (C) Binary and (D) Ternary polymer-membrane systems.

### 3.3 Lipid Sorting and Co-localization with defects

To this end, we first calculate the total surface areas attributed to the two lipid types: POPE and POPG in a given frame. We then determine the fraction of each surface area co-localized with deep defects,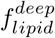. The calculation is performed for the two leaflets, *L*0 and *L*1 separately over the equilibrated trajectories of last 500 ns for both polymer-membrane systems to generate distributions, 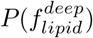. In presence of the binary polymer (Fig. 4C) the intra-leaflet distributions, 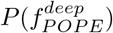 and 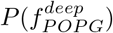 significantly overlap. This indicates that there is no preference for a particular lipid type to form defects. However, we observe significant differences in distributions between the two leaflets. Owing to abundance of deep defect sites of varying sizes in the proximal *L*0 leaflet, the defect area fractions corresponding to the two lipid types are significantly higher, indicated by the broad distributions compared to more sharp distributions observed in the lower *L*1 leaflet.

Similarly in presence of ternary polymer, the distributions of defect area fractions, 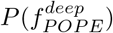 and 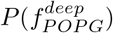 of distal leaflet are also overlapping (Fig. 4D). However, the proximal leaflet *L*0 is characterized by significantly higher defect area fraction corresponding to POPG lipids than POPE lipids. The deep defects in *L*0 are thus co-localized with POPG under the influence of ternary polymer. This apparent co-localization is a direct manifestation of POPG clustering in vicinity of ternary polymer insertion site. Incidentally this insertion site being co-localized with a single large deep defect results in POPG enriched defect site. This is also evident from Fig. 4B, which illustrates crowding of POPG molecules surrounding the ternary polymer embedded deep defect site.

## 4. Discussion

In the present work we elucidate the role of interfacial packing defects in recruiting antimicrobial polymers into bacterial membrane. The action of antimicrobial polymers enhances both the number and the size of lipid packing defect sites upon approaching the proximal leaflet of bacterial membrane. The hydrophobic residues of both binary and ternary polymers acting as packing defect sensors, facilitate discrete insertion events into available deep defect sites, thereby resulting in partitioning of these polymers into membrane milieu. On other hand, the presence of polar residues in ternary polymer tend to occupy shallow defect sites close to the headgroup-solvent interface. The competitive tendency of hydrophobic and polar residues to occupy defect sites of varying depths along with the enhanced hydrogen bonding network induce a more folded conformation of ternary polymer, resulting in partition mechanism different from that of the binary polymer, which acquires facially amphiphilic conformation upon partitioning.

The presence of charged cationic functional groups within the folded conformation of ternary polymer may exhibit local charge density. Such cationic charge density in different antimicrobial and cell penetrating peptides is known to induce sorting of anionic POPG lipids[18, 29, 31, 39, 40, 41]. Similar clustering of POPG lipids is observed in the neighbourhood of ternary polymer insertion site leading to apparent co-localization with lipid packing defects. Although conical lipids with relatively small head groups compared to lipid tail cross section, such as POPE, are known to promote defect formation, no preferential bias towards occupying defect sites is observed for either POPE or POPG lipids. A similar observation was noted for DOPC-DOG membrane, where presence of conical DOG lipids enhanced lipid packing defects but did not co-localize with defect sites [23].The presence of conical lipids thus introduce defects randomly distributed all over the membrane surface.

Several studies including the present one, have provided useful insight into mechanisms of partitioning being regulated by sensing of lipid packing defects[22, 23, 24, 25, 26]. Recently, it has been concluded that viral peptide entry into host cell membranes follow a similar mechanism by sensing membrane topography[27, 28]. MD simulations of the viral peptide in presence of model POPC bilayers revealed the presence of lipid packing defect sites, which act as transient binding spots facilitating peptide partitioning. Further, presence of cholesterol significantly altered membrane topography in terms of reduced lipid packing defects, thereby mitigating viral peptide entry into host cell membranes. Although bacterial membrane is different from mammalian cell membrane in terms of lipid type and composition, packing defects are inherently present across different biomembranes. As a result, partitioning of different classes of membrane active agents, like antimicrobial peptides / polymers or viral peptides follow a general mechanism through sensing of such lipid packing defects.

## 5. Conclusion

In the present work, we provide thorough insight into the role of lipid packing defects upon interaction of biomimetic antimicrobial polymers with model bacterial membranes. The partitioning dynamics of ternary polymers composed of charged cationic, hydrophobic and neutral polar groups is compared with that of binary polymer, the latter being devoid of any polar groups. The presence of hydrophobic groups in either polymers enhances deep defect sites on the proximal leaflet and facilitate insertion and subsequent partitioning of the polymers. The shallow defect sites, on the other hand, are moderately enhanced only in presence of the ternary polymer, owing to intermittent residence of polar groups close to the lipid-water interface. The competitive tendency of hydrophobic and polar residues of ternary polymer to occupy deep and shallow defect sites, respectively, accompanied by strong hydrogen bonding network induce a more folded conformation, resulting in partition mechanism different from that of the binary polymer, which acquires a facially amphiphilic conformation upon partitioning. Here we provide conclusive evidence that insertion of antimicrobial polymers in bacterial membrane is preceded by sensing of interfacial lipid packing defects, a mechanism similar to other reported membrane active agents.

## Acknowledgement

All the simulations in this work have been carried out on clusters Annapurna and Nandadevi at The Institute of Mathematical Sciences, Chennai, India.

